# From wings to tails: a general framework for inferring dispersal potential from morphology

**DOI:** 10.64898/2026.07.16.739000

**Authors:** Rachel M. Aitchison, Wade J. VanderWright, Jay H. Matsushiba, Nicholas K. Dulvy

## Abstract

Dispersal shapes gene flow, range dynamics, and extinction risk, yet direct measures of dispersal are unavailable for most taxa. Here, we test whether the aspect ratio of propulsive surfaces, an established proxy in birds, provides a general morphological basis for inferring dispersal potential. Using sharks as a model taxa, we find caudal fin aspect ratio was positively associated with swimming speed and geographic range size. Applying this proxy across 452 species, we find no evidence of a latitudinal gradient in dispersal potential, in contrast to dispersal patterns reported for birds, due to the lack of a direct effect of environmental temperature on the geographic distribution of caudal fin aspect ratio. Higher dispersal potential was associated with pelagic habitats, larger body size and greater oxygen uptake capacity. Because aspect ratios can be measured from museum specimens and illustrations, this approach could enable comparisons of dispersal potential across air- and water-dwelling vertebrates.

## INTRODUCTION

Trait-based approaches offer a scalable means of estimating ecological functions and responses to threats that are otherwise difficult to measure directly, especially in data-poor lineages(*1–3*). Ecological function can be defined in terms of rates (e.g., rates of nitrogen deposition) and states that are indirectly related to ecological processes through rates (e.g., biomass is tied to productivity)(*2*). Understanding ecological function has historically relied on ‘big-data’ analyses that offer generality (e.g., across plants or coral reefs), but these studies have been critiqued for the limited attention paid to whether traits available in public databases are genuinely related to ecological function(*2*). The alternative is painstaking ecological research to establish causal connections to ecological rates(*2*, *3*). Morphological traits, however, are well-suited candidates for estimating function and difficult-to-measure ecological processes.

Dispersal, or the movement of individuals through space, is a key functional process that influences ecological processes including population connectivity and viability(*4*, *5*), nutrient transfer(*6*, *7*), and community assembly(*7*, *8*), as well as the evolutionary process of speciation(*9*). We have little comparative understanding of the drivers of dispersal, mainly because there are no lineage-wide estimates of dispersal potential except for birds. This is because dispersal remains difficult to systematically quantify in a representative manner across large numbers of species. The challenge is that methods for quantifying dispersal, such as mark-recapture and electronic tagging, can be expensive and time-consuming to apply across all species in a lineage and the numbers of animals tagged rarely represents population-scale processes. For example, nearly 100 years of mark-recapture efforts in Britain resulted in natal and breeding dispersal data for only 75 species of bird(*10*). In the marine realm, acoustic telemetry has been deployed on only 70 species of shark out of 530+(*11*, *12*), and satellite tags have only been applied to a few dozen large-bodied shark species(*13–15*). Consequently, there are significant taxonomic and geographic biases in tracking data, with most studies limited to the most charismatic species and most developed nations of Europe and North America, followed by Australia and South Africa, with much of the Global South lacking movement data(*13*, *16–18*).

Further, there is a conservation imperative to quantify and compare dispersal across whole lineages. First, dispersal underlies recovery potential from threats such as overfishing(*6*), climate change, and habitat loss(*19*). Second, the Kunming-Montreal Global Biodiversity framework targets specify the need to halt and reverse the loss of ecosystem functions and services, including ecological integrity and connectivity (e.g., Targets 3, 4(*20*)). Third, the Convention on Migratory Species requires the identification of at-risk migratory species for protection(*21*).

In birds, wing morphology provides a functional trait for estimating dispersal potential(*22*). Specifically, the aspect ratio of wing span and width, also known as Hand-Wing Index, can be readily calculated from museum specimens(*19*) and demonstrably predicts flight efficiency(*23*) as well as geographic range size(*19*, *24*). Avian wing shape, and dispersal potential, is correlated with seasonality, body size, as well as migratory and territorial behaviour(*19*). Thus, avian Hand-Wing Index provides an established example of how the shape of a propulsive surface can be used to infer dispersal ability across an entire lineage. Since high aspect ratio wings and fins are associated with sustained locomotion in air and water(*25*), a similar relationship between fins and dispersal may hold for aquatic organisms. Although fin aspect ratio is a well-studied correlate of swimming performance in fishes(*26–29*), it has not been evaluated as a lineage-wide measure of dispersal potential.

Sharks are a diverse basal lineage of jawed vertebrates(*30*) that represent an independent evolutionary origin of propulsion (i.e., tail fins instead of wings), providing a test case for whether morphology can predict dispersal in the aquatic realm as has been done for the aerial realm. Similar to birds, sharks fill a wide range of ecological roles(*6*), occupy a variety of habitats(*31*), and are globally distributed(*6*, *30*, *31*). Also, despite substantial variation in body size and shape, all sharks rely on caudal fins as the main mode of propulsion. A morphological analogue to Hand-Wing Index has been established for sharks - caudal fin aspect ratio (CFAR)(*32*).

Species with higher CFAR values have more symmetric homocercal caudal fins (e.g., White Shark, *Carcharodon carcharias*) whereas species with lower CFAR values have asymmetric heterocercal caudal fins (e.g., Nurse Shark, *Ginglymostoma cirratum*). Species with higher CFAR values (taller with lower surface area) generate greater thrust(*33*), swim faster(*28*), and have longer gill slits(*29*), which are positively related to gill surface area(*34*), compared to sharks with lower CFAR values. CFAR can be measured from anatomically accurate illustrations in field guides(*29*) and validated with museum specimens, and it is easy to reliably measure across many species. CFAR ranges from 0.25 in the tide-pool dwelling Epaulette Shark (*Hemiscyllium ocellatum*) to 5.29 in the ocean-transiting Longfin Mako (*Isurus paucus*)(*29*). Whether CFAR is a reliable measure of dispersal in sharks has not been tested, nor have the ecological and evolutionary factors that shape variation in dispersal potential across the lineage. If propulsor aspect ratio tracks dispersal in both birds and sharks, two disparate vertebrate lineages, this would support aspect ratio as a general morphological basis for inferring dispersal potential in fluid environments. Such a proxy would also make it possible to ask whether broad-scale patterns in dispersal, such as latitudinal gradients, are shared between aerial and aquatic systems or reflect the distinct ecological adaptations of each.

Here, we develop and apply caudal fin aspect ratio as a morphological proxy for dispersal potential, using sharks as a model lineage, and test the ecological and evolutionary factors that shape dispersal potential. First, we assess the taxonomic and geographic coverage of existing tracking data to evaluate the suitability for lineage-wide estimates of dispersal. Second, we test CFAR as a proxy for dispersal potential against the swimming speed of free-living sharks and against geographic range. Third, we describe present-day phylogenetic and geographic patterns in dispersal potential, including whether the latitudinal gradient reported for birds is also found in sharks. Finally, we test the hypothesized climatic, ecological, and morphological drivers (Table 1) that may underlie these phylogenetic and geographic patterns.

**Table 1.** Proposed hypotheses of the causal relationships between variables. Relationship between each variable, direction (positive or negative), proposed explanation, and references to support each relationship.

| Relationship | Direction | Proposed Explanation | References |
| --- | --- | --- | --- |
| Average Temperature drives Maximum Body Size | Negative | Greater average temperatures drive smaller body sizes (temperature size rule) through metabolic rate. | Atkinson (1994)(36) |
| Maximum Body Size drives Gill Slit Height | Positive | Larger body size drives gill slit height due to greater metabolic demand through gill surface area. | Bigman et al. (2018)(32), Pauly (2021)(73), VanderWright et al. (2024)(29) |
| Maximum Body Size drives CFAR | Positive | Indirectly through gill slit height (gill surface area) due to metabolic demand, and directly as larger sharks have greater foraging areas. Sharks likely originated as smaller-bodied and benthic. | VanderWright et al. (2024)(29), Sternes et al. (2024)(40) |
| Water Column Position drives Maximum Body Size | Positive (Pelagic Body Size > Benthic Body Size) | Pelagic species should have larger maximum body sizes due to selective pressure in the more homogeneous environment of the pelagic zone (compared to the benthic zone) with the expansion of sharks into the pelagic zone (from a likely benthic ancestral state). | Compagno (1990)(31), Friedman et al. (2020)(74), Sternes et al. (2024)(40) |
| Water Column Position drives Gill Slit Height | Positive (Pelagic Gill Slit Height > Benthic Gill Slit Height) | Water column position drives gill slit height due to differences in the metabolic demand associated with each position (pelagic species have greater metabolic demand). | Bigman et al. (2018)(32) |
| Water Column Position drives CFAR | Positive (Pelagic CFAR > Benthic CFAR) | Pelagic species should have a higher CFAR than benthic species due to the need to move constantly (especially ram ventilators) and to improve swimming efficiency by reducing drag. | Compagno (1990)(31), Bigman et al. (2018)(32), Grady et al. (2022)(75), Iliou et al. (2023) (28), VanderWright et al (2024)(29), Sternes et al. (2024)(40) |
| Gill Slit Height drives CFAR | Positive | Gill slit height drives CFAR due to lifestyle and | Rubio-Gracia et al. (2020)(76), Fu et al. |
|  |  | metabolic demand. A co-evolution of metabolic “capacity” (gills) and “demand” (tails) is likely, however, increased capacity would permit increased demand. | (2022)(77), VanderWright et al. (2024)(29) |

## RESULTS

### Patterns in dispersal data from tagging

Electronic tracking studies were strongly non-random across evolutionary, ecological and conservation axes (Fig. S1, Table S1). There is greater focus on tracking large-bodied pelagic species (*χ*^2^ = 51.036, df = 2, p < 0.01), and tagging is especially phylogenetically biased toward requiem sharks (Carcharhinidae; *χ*^2^ = 177.02, p < 0.01; Monte Carlo simulation, 5,000 replicates). There is also a tendency to tag species at lower threatened IUCN categories (e.g., Vulnerable versus Critically Endangered; *χ*^2^ = 25.767, df = 5, p < 0.01), presumably because they are more widespread and abundant. Thus, current electronic tracking data cannot yet be used to infer general patterns and drivers of dispersal without additional work to expand phylogenetic and ecological coverage.

### CFAR as a proxy for dispersal potential

Caudal fin aspect ratio (CFAR) ranges from 0.39 to 5.29(*28*) (Fig. 1a). is positively related to swimming speed in wild sharks (n = 23 Fig. 1b) and geographic range size (n = 452; Fig. 1c) such that species with greater CFAR values have faster swimming speeds (*β* = 0.15, 95% Credible Interval (CI) = -0.18–0.43; Table S2) and larger geographic ranges (*β* = 13.17, 95% CI = 10.16–16.20; Table S2).

**Figure 1.**
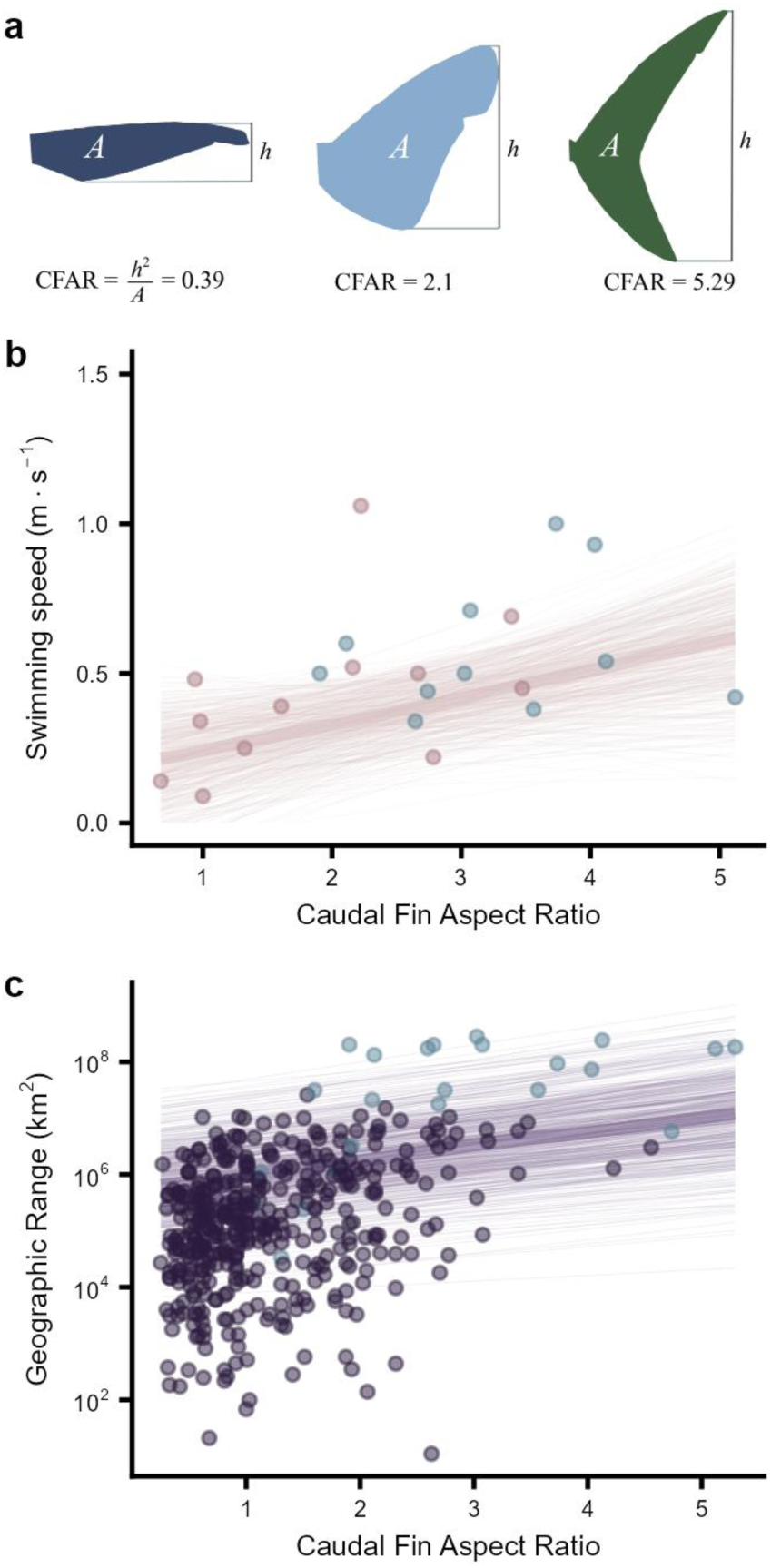
Caudal fin aspect ratio (CFAR) as a measure of dispersal potential. (a) Variation in CFAR with three examples (low CFAR: Longfin Catshark, *Apristurus herklotsi*; medium CFAR: Prickly Dogfish, *Oxynotus bruniensis*; high CFAR: Longfin Mako, *Isurus paucus*). Black boxes represent the caudal fin height (*h*) and tail shading represents surface area (*A*). (b) Swimming speed in the wild as a function of CFAR for 23 species, modified from Iliou et al.(*28*). (c) Geographic range size as a function of CFAR for 452 species. Lines are 500 random draws of conditional fit from the posterior distribution of each model with the median draw line emphasized. Pelagic species are emphasized in blue.

### Phylogenetic patterns in dispersal potential

Greater dispersal potential is coincident with larger body sizes, greater oxygen uptake potential (i.e., longer gill slits), and in pelagic species (Fig. 2). The intercorrelation between these three factors is most notable in some ground sharks (Carcharhiniformes) and the mackerel sharks (Lamniformes) which have the greatest CFAR and thus dispersal potential suggesting coevolution of body size, oxygen uptake and caudal fin aspect ratio. CFAR is lowest in the bramble sharks (Echinorhiniformes), carpet sharks (Orectolobiformes with the exception of the Whale Shark *Rhincodon typus* which has a CFAR of 5.12), cow sharks and frilled sharks (Hexanchiformes), and sawsharks (Pristiophoriformes).

**Figure 2.**
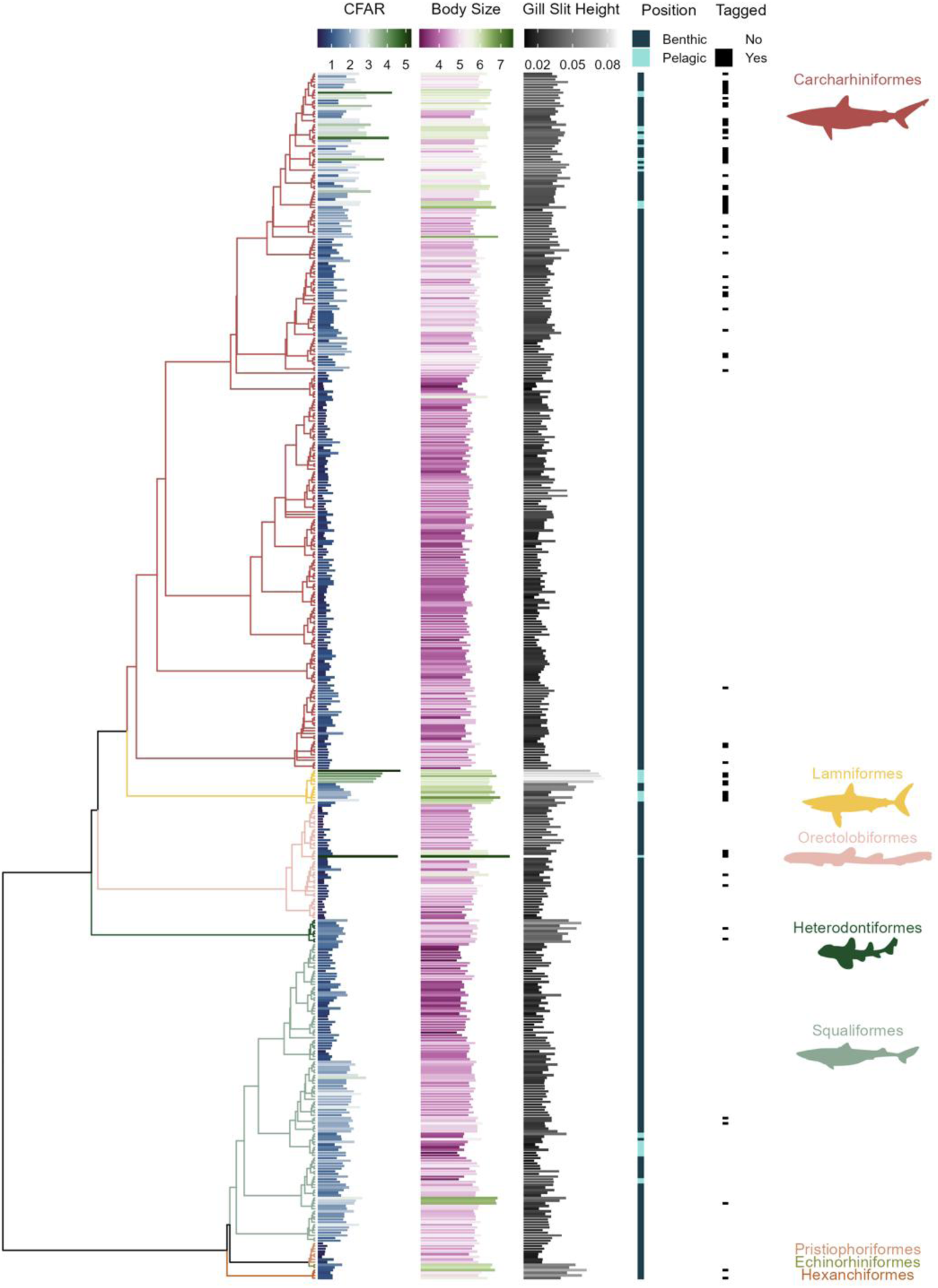
Phylogenetic patterns in caudal fin aspect ratio (CFAR) and associated traits. The eight orders represented are labeled, and each column indicates trait values. From left to right: caudal fin aspect ratio (CFAR), blue colors indicate lower values and green colors indicate higher values; maximum body size (logged, base 10), pink colors indicate smaller species and green colors indicate larger species; oxygen uptake potential (gill slit height, % total length), dark grey colors indicate lower oxygen uptake potential and light grey colors indicate higher oxygen uptake potential; water column position, dark blue indicates benthic and light blue indicates pelagic; tagged, black indicates species with tracking data and white indicates species without tracking data. Silhouettes are from phylopic.org (open license, CC0) and are unchanged except coloration.

### Geographic patterns in dispersal potential

Shark species richness is greatest in the tropics (Fig. 3a), yet there is no latitudinal gradient in mean CFAR per cell (Fig. 3b). Variability in CFAR has a slight latitudinal gradient where variability is lower towards low latitudes (<30°) and greater towards middle latitudes (30°–60°; Fig. 3c). Instead, there is a strong coastal-pelagic gradient in CFAR where mean CFAR is lower along the coasts and islands, and higher offshore in all ocean basins (Fig. 4a). When considering all sharks, the coastal-pelagic gradient in mean CFAR (Fig. 4a) is slightly obscured by the presence of offshore and seamount deepwater sharks, resulting in lower CFAR than expected in the open ocean (Figs. 4c and g). We find a similar coastal-pelagic pattern in the variability in CFAR per cell (Fig. 4b) where variability is greater just offshore than along the coast, though the gradient is much less pronounced than mean CFAR. We also find that the distribution of mean logged maximum body size is nearly identical to the distribution of mean CFAR (Figs. 4a and S8).

**Figure 3.**
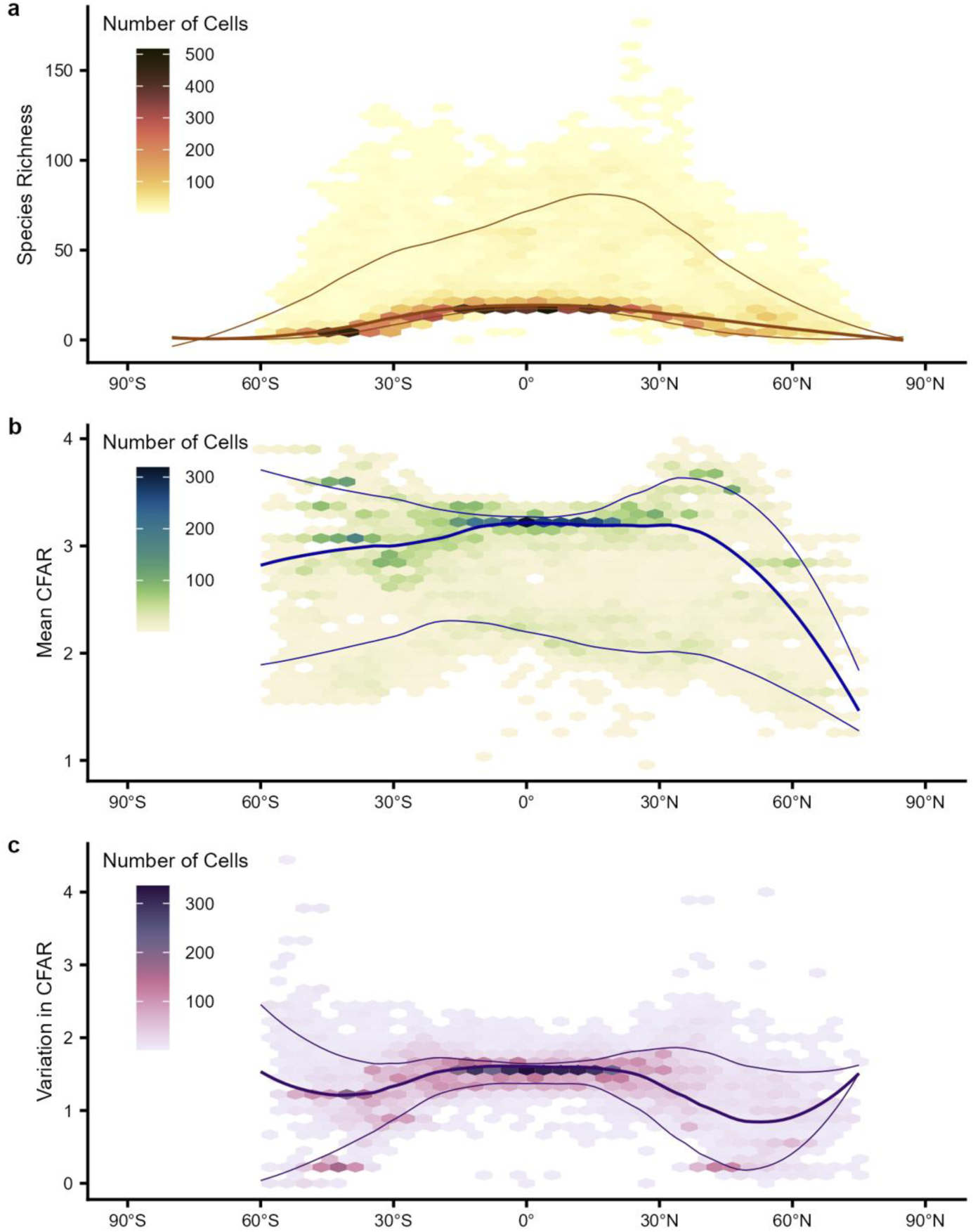
Latitudinal patterns of shark diversity and mean and variance in dispersal potential (CFAR). (a) Global shark species richness (n = 452 species). (b) Mean CFAR, weighted by species richness. (c) Variation in CFAR, weighted by species richness. Loess lines represent the 90th, 50th, and 10th centiles with a span of 0.5.

**Figure 4.**
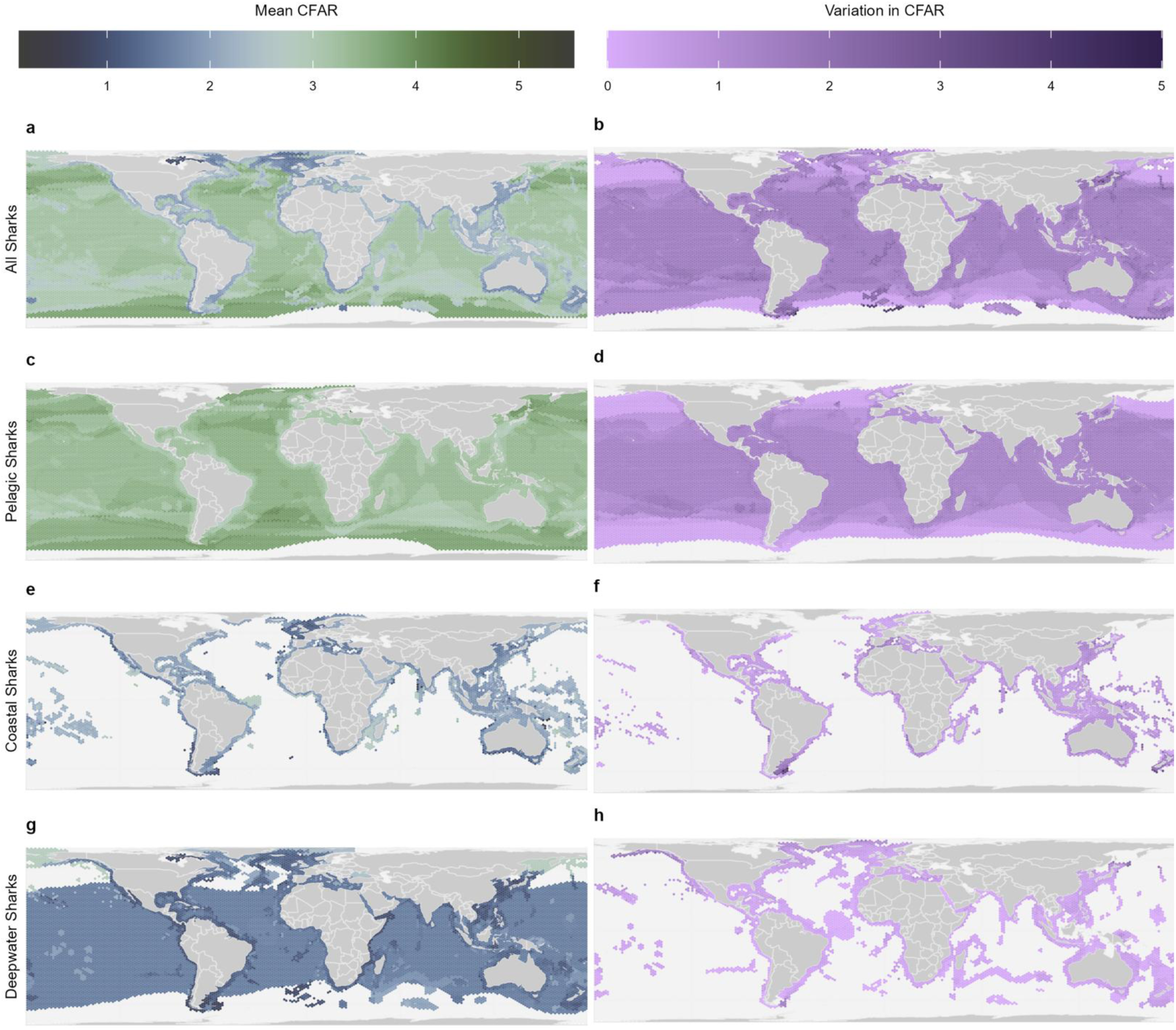
Global patterns in dispersal potential (caudal fin aspect ratio, CFAR). Mean CFAR per cell for (a) all sharks (n = 452), (c) pelagic species (n = 25), (e) coastal species (n = 198), and (g) deepwater species (n = 229). Blue colors indicate lower CFAR and green colors indicate higher CFAR. Variability in CFAR for (b) all sharks, (d) pelagic species, (f) coastal species, and (h) deepwater species. Lighter purple colors indicate lower variability and darker purple colors indicate greater variability. Each cell represents approximately 23,322 km^2^.

### Causal modeling of dispersal potential

In order to understand the drivers of the coastal-pelagic pattern in CFAR given the clear correlation in traits (Fig. 2), we evaluated four causal models in `phylopath` (Figs. S5a, S6a, and 5) based on hypotheses in the literature (Table 1). Our final model passed the d-separation test, meaning all paths between variables are conditionally independent thus removing potential confounds (Fig. 5, Table S11)(*33*). Dispersal potential is driven by metabolically-relevant morphological traits and ecological factors with no direct climatic effect of temperature, which explains the absence of a latitudinal gradient in CFAR (Fig. 4a,b; Fig. 5). Maximum body size had a positive direct effect on CFAR (*β* = 0.17) with oxygen uptake potential (relative gill slit height) acting as a strong mediator. Maximum body size had a positive effect on gill slit height (*β* = 0.38), which in-turn had a positive effect on CFAR (*β* = 0.21). Water column position had strong positive effects on maximum body size (*β* = 0.33), gill slit height (*β* = 0.41), and CFAR (*β* = 0.51). Water column position is also strongly associated with habitat use (coastal or oceanic, Figs. S4 and S7). Finally, average temperature had a strong negative effect on maximum body size (*β* = -0.38) as expected from metabolic theory and the temperature-size rule(34). The effect of maximum body size, both direct and indirect, aligns with the coastal-pelagic gradient, such that smaller species with lower dispersal potential are found in coastal ecosystems, whereas the correlated larger body sizes, greater uptake potential and CFAR coincides with the occupation of the pelagic realm (Figs. 4 and 5).

**Figure 5.**
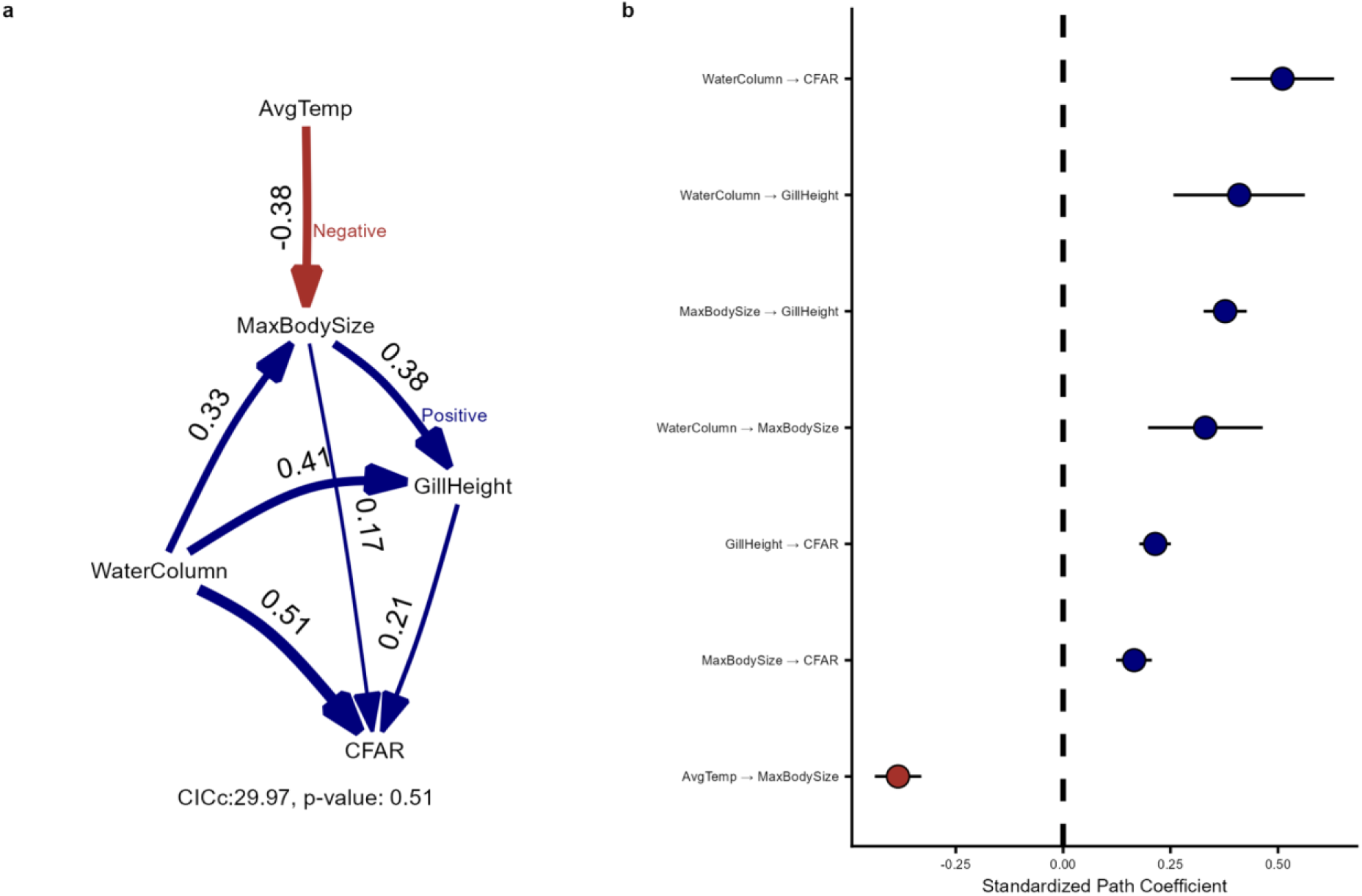
Phylogenetic path analysis of the drivers of shark dispersal potential (CFAR). (a) The best supported model and coefficients showing the direction and strength of effects. Arrow thickness is proportional to coefficient strength and color indicates direction: negative (red, e.g. AvgTemp to MaxBodySize) and positive (blue, e.g. MaxBodySize to GillSlitHeight). The associated C-information criterion (CICc) value and model *p*-value (a significant *p*-value indicates rejection in phylopath) are included (see also Table S11). (b) Standardized path coefficients with standard error intervals for each path.

### Bayesian models of dispersal potential informed by causal modelling

Our Bayesian models confirm metabolically-relevant morphological traits and ecological factors drive global patterns in CFAR (Fig. 6). Maximum body size (while accounting for water column position, Table 2) and gill slit height (while accounting for maximum body size and water column position, Table 2) had positive effects on CFAR (*β* = 0.15 and 0.14, respectively). Thus, larger-bodied species with greater oxygen uptake capacity (relative gill slit height) have greater dispersal potential than smaller-bodied species. Average temperature had a negative effect on CFAR (*β* = -0.11), such that species in warmer waters have lower dispersal potential. There was a small difference in dispersal potential between benthic and pelagic species. Specifically, the estimated marginal means for the CFAR of benthic species was lower than pelagic species (*β* = 1.22 versus *β* = 1.93, respectively). The results of our Bayesian analysis further explain the coastal-pelagic gradient by illustrating the absence of a latitudinal gradient is due to the lack of a strong direct temperature effect on CFAR (Figs. 4 and 6).

**Figure 6.**
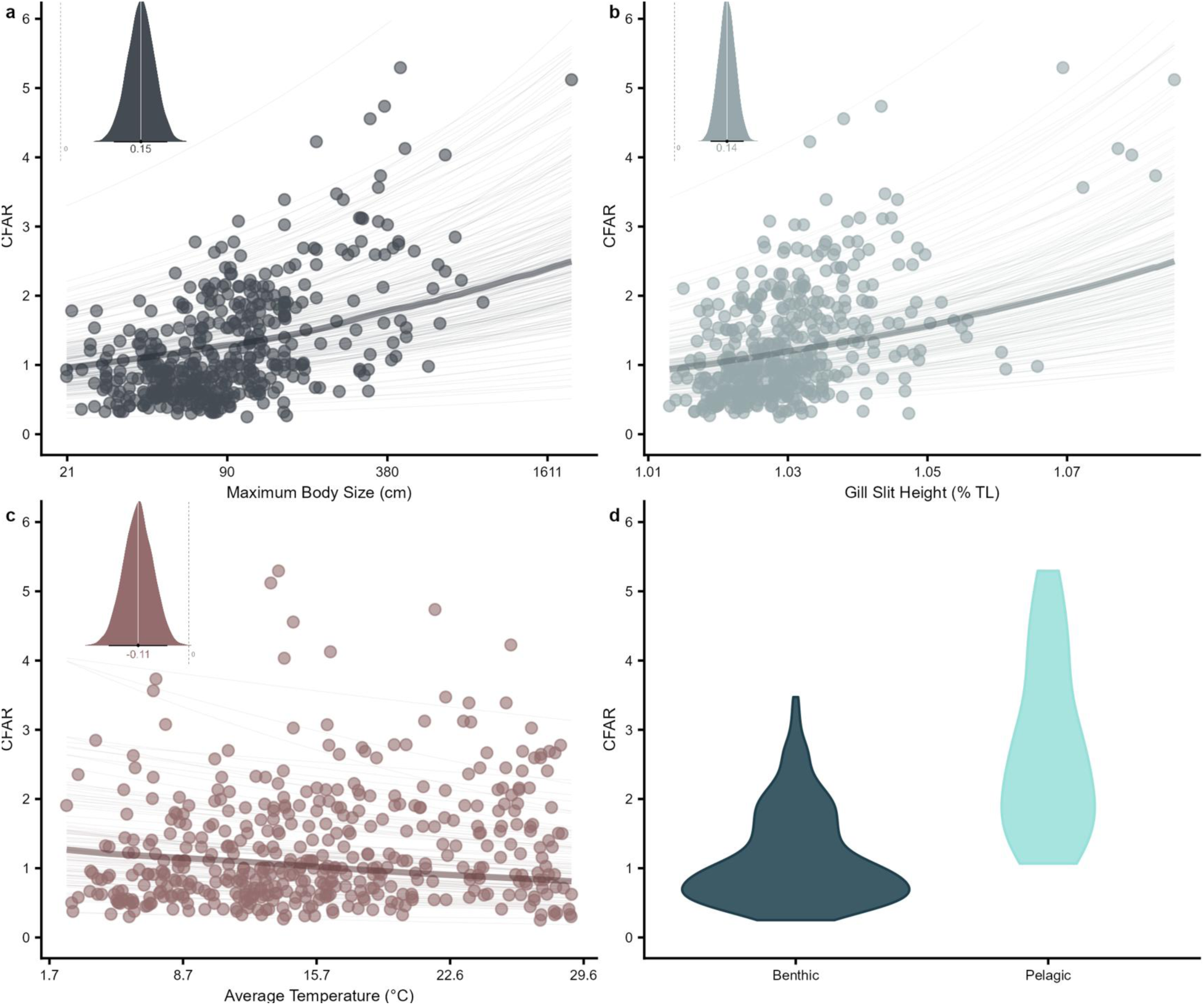
Bayesian analysis of the drivers of shark dispersal potential (CFAR). (a) Maximum body size (logged). (b) Oxygen uptake potential (gill slit height, % total Length). (c) Average temperature (°C). (d) Water column position. In panels a-c, the posteriors from each model are inset showing the coefficient value and departure from zero. Lines represent 500 random draws of conditional fit from the posterior distribution with the median draw emphasized.

**Table 2.** Adjustment sets for Bayesian models. Adjustment sets for determining the total effect of each variable on CFAR from DAGitty with the variable of interest in bold.

| Variable | Adjustment formula |
| --- | --- |
| Average Temperature | CFAR ~ **Average Temperature** |
| Maximum Body Size | CFAR ~ **Maximum Body Size** + Water Column Position |
| Gill Slit Height | CFAR ~ **Gill Slit Height** + Maximum Body Size + Water Column Position |
| Water Column Position | CFAR ~ **Water Column Position** |

## DISCUSSION

Our results indicate caudal fin aspect ratio (CFAR) can serve as a morphological proxy for dispersal potential in sharks, enabling lineage-wide inferences that are not yet feasible from tracking data. Across 452 species, we found no evidence of a latitudinal gradient in dispersal potential, in contrast to birds. Instead, dispersal potential was associated primarily with ecological specialization, particularly pelagic habitat use, and with larger body size and greater oxygen uptake capacity. Overall, the use of aspect ratios of the propulsive surface to track dispersal potential in two disparate lineages (birds and sharks) supports the use of aspect ratios as a general basis for comparing dispersal potential across flying and swimming vertebrates. This approach may also extend to extinct taxa, such as ichthyosaurs, and migratory flying insects.

Applying this dispersal potential framework to sharks reveals null morphological estimates of how dispersal potential is shaped by species ecology, rather than latitude, highlighting how functional morphology can be used to uncover the mechanisms driving movement ability at broad scales. More broadly, trait-based approaches provide a cost-effective tool for filling data gaps without sampling rare or threatened taxa. Next, we discuss: (i), why is there no latitudinal gradient in shark dispersal?, (ii) why is there a coastal-pelagic gradient in shark dispersal?, (iii) the evolution of shark tail shape, (iv) conservation implications, and (v) future directions.

Birds exhibit a marked latitudinal gradient in dispersal(*19*) but our analysis indicates sharks do not: we now know this only from these trait-based analysis rather from tracking studies. Bird wing shape is closely related to migration behaviour and seasonality (breeding range temperature variability)(*19*). As a consequence of intense, short seasonal production at high latitudes, it is evolutionary advantageous to migrate long distances due to the payoff of enhanced feeding and breeding opportunities, especially in the Arctic(*19*). This is also seen in diadromy in fishes which is thought to have evolved at higher latitude to allow smaller freshwater life stages to take advantage of seasonally high marine productivity and mid- to high latitudes fishes(*37*). By contrast, shark dispersal is not directly related to environmental temperature explaining the absence of a general latitudinal gradient in CFAR. Interestingly, poleward seasonal migrations of sharks are known in some places, notably along the eastern seaboard of the USA(*38*, *39*). This serves to remind us that the detail and complexities of dispersal of individual species depended on the interaction of behaviour and oceanography at smaller scales. As such, trait-based approaches thus provide a first approximation of dispersal potential across species complemented by spatially and temporally more detailed electronic tracking studies.

The coastal-pelagic gradient in shark dispersal may be a result of the evolution and habitat shift by select lineages of sharks into the pelagic ocean, notably some requiem sharks and mackerel sharks(*31*). Sharks are hypothesized to have evolved from small bottom-dwelling coastal species, with subsequent expansion into the deepsea and pelagic ocean(*31*, *40*). Our analysis suggests the invasion of the pelagic ocean likely involved the concerted evolution of larger body size, increased oxygen uptake capacity, elongated pectoral fins(*40*), and greater caudal fin aspect ratios, resulting from the evolution of more symmetrical (homocercal) tails (Fig. 2). These changes led to increased lift and thrust(*33*), swimming speeds, and cruising capacity(*28*), enabling populations to sustain connectivity between discrete habitats. Not only do pelagic species range widely along frontal systems and gyres, tracking ephemeral production(*41*), but many species dive deep to intercept prey in the oxygen minimum zone or deep scattering layer(*42*), and give live birth in coastal areas(*43*, *44*). All three behaviours require significant dispersal and metabolic capacity, enabled by greater oxygen uptake capacity and larger body sizes.

In the current biodiversity crisis, where over one-third of sharks and rays are threatened with extinction(*45*), the application of causal inference of dispersal potential can support effective species and area-based management strategies(*30*). The phylogenetic and geographic patterns of dispersal potential outlined here can be used to identify regions where species have limited dispersal potential to inform the strategic placement of spatial protections to ensure connectivity, such as marine protected areas (MPAs)(*46*) and Important Shark and Ray Areas (ISRAs)(*47*). Species with low CFAR are potentially more sensitive to habitat loss and overfishing, and the phylogenetic and geographic distribution of mean CFAR reveal where these threats might be most acutely realized. Universally, we find coastal areas have lower CFAR and thus lower dispersal potential. In coastal reef ecosystems, for example, the overfishing of larger-bodied shark species has led to changes in community composition(*6*), resulting in a decrease in mean CFAR, reduced dispersal potential, and loss of ecological connectivity between reefs. Regions with low mean CFAR overlap with hotspots of threatened endemic species (e.g., southeastern Australia, South Africa, and southeastern China)(*18*, *30*), supporting the connection between low dispersal potential and increased likelihood of speciation and elevated extinction risk. In addition, inferred dispersal potential through CFAR can be used as part of a trait-based approach to infer the extinction risk of data-poor newly described species.

We encourage additional electronic tracking analyses to validate the connection between morphology and dispersal, such as the ontogenetic expansion of dispersal potential as tail shape changes with body size. We hypothesize that CFAR will be strongly related to dispersal kernel distances and genetic connectivity (isolation-by-distance), further strengthening our understanding of CFAR as a measure of the ecological functionality of dispersal potential(*2*). We could not include the rays, (Superorder Batoidea), which are the sister taxon to sharks, due to the lack of an established morphological trait that connects ray morphology to metabolism and swimming speed. We hypothesize that ray fin morphology will also explain variation in swimming speed, dispersal distance, genetic connectivity and geographic range size(*48*, *49*). Expanding our understanding of the bridge between metabolic morphology, life histories, and population growth rate would allow us to compare dispersal potential and susceptibility to threats for both sharks and rays, encompassing all elasmobranchs (1,200+ species)(*50*, *51*). We conclude that trait-based measures of dispersal hold great promise for generalizing and understanding the evolution and ecology of dispersal potential across vertebrates in the air and aquatic realms potentially informing the next wave of spatial protections focusing beyond patterns of species richness, endemicity and evolutionary distinctiveness but also on ecological functions, particularly ecological connectivity.

## METHODS

In order to understand the geographic, climatic, ecological, and morphological drivers of caudal fin aspect ratio (CFAR), we first describe how CFAR is measured, and how it connects to dispersal potential. We then describe how we obtained climatic (average temperature at depth, average dissolved oxygen at depth), ecological (habitat use, water column position), morphological (maximum body size, gill slit height), and phylogenetic data. Finally, we explain how we mapped the global distribution of CFAR and illustrate our causal inference approach and statistical analyses.

### Calculating caudal fin aspect ratio

Caudal fin aspect ratio (CFAR) measurements were taken from the field guide *Sharks of the World: A Complete Guide*(*11*), which has been used in previous studies(*28*, *29*, *40*). CFAR was calculated as CFAR *= h² / A*(*28*, *29*) where *h* is the height of the caudal fin and *A* is the surface area of the caudal fin measured in ImageJ (Fig. 1A).

### Evaluating bias in tracking data

We first tested whether the electronic tagging data available for sharks(*13*, *16*) were biased with respect to ecology, taxonomy and conservation status with chi-square tests of independence between whether a species has been tagged or not (“Y” or “N”) and water column position (“Benthic” or “Pelagic”), taxonomic family, and IUCN Red List category. To assess bias in tracking by taxonomic family, we estimated significant bias by estimating p-values via Monte Carlo simulation with 5,000 replicates.

### Evaluating swimming speed and geographic range

To evaluate CFAR as a measure of dispersal potential, we tested the hypothesis that CFAR is positively related to (i) instantaneous swimming speeds and (ii) global geographic range using Bayesian generalized linear multilevel models using the brms package(*52*), with phylogeny as a random effect. Swimming speed data (m s^-1^) were obtained from previous studies on sharks(*28*, *53*) and species range sizes (km^2^) were obtained from the IUCN Red List and Dulvy et al. 2021(*45*, *54*). All models were fitted with four chains each of 3,000 iterations with 1,000 warm-up iterations (4,000 iterations total). Model convergence was assessed by ensuring R-hat values were equal to one and effective sample size (ESS) were greater than 1,000. All analyses were performed using R (v 4.6.1).

### Extracting climatic, ecological, and morphological variables

We extracted average temperature-at-depth and dissolved oxygen-at-depth for each species, based on their geographic ranges (see “Mapping global patterns of dispersal potential”) and depth ranges. We extracted the upper and lower depths for species through the IUCN Red List API using the iucnredlist package(*55*). We used the National Oceanographic Administration (NOAA) World Ocean Atlas, as it provides values for climatic variables at regular points across the sea surface and at standard depths(*56*). Using the sf and terra R packages(*57*, *58*), we queried the climatic data, returning values from cells that intersected with species range, by area and depth. From these values, we calculated the annual mean temperature and average dissolved oxygen values for each species.

Gill slit height was measured from illustrations in *Sharks of the World* and expressed as a proportion of total length(*11*, *29*). We use gill slit height as a measure of oxygen uptake potential as it is strongly positively related to gill surface area(*34*) and gill surface area data are not available for most shark species(*34*, *59*). Maximum body size (total length in cm) was obtained from the International Union for the Conservation of Nature (IUCN) Red List and Dulvy et al.(*45*). Maximum body size was log-transformed and all quantitative (continuous) variables, including logged maximum body size, were centered and scaled by subtracting the mean and dividing by one standard deviation.

Water column position (benthic, n = 254; benthopelagic, n = 202; pelagic, n = 45) was obtained from Pimiento et al.(*60*) and the Habitat & Ecology narratives from the IUCN Red List assessments. Habitat use (coastal, n = 229; deepwater, n = 289; pelagic [oceanic], n = 28) was obtained from the IUCN Red List and Dulvy et al.(*45*). Our causal modeling framework only supports binary categorical variables so we used phylogenetic generalized least squared (PGLS) models and compared Akaike Information Criterion (AIC) values to reduce the number of categories for water column position (benthic, benthopelagic, and pelagic) and habitat use (coastal, deepwater, and pelagic [oceanic]). We determined “benthopelagic” should be merged with “benthic” (Fig. S2, Table S3) and “deepwater” should be merged with “coastal” (Fig. S3, Table S4).

After reducing our categorical variables (water column position and habitat use) to a binary, we inspected species classifications (e.g., the Scalloped Hammerhead, *Sphyrna lewini*, was classified as ‘Pelagic’ and ‘Oceanic’) to ensure category assignments were consistent and ecologically sound. Some species classifications, particularly water column position, were manually changed to better reflect the IUCN Red List Habitat & Ecology narratives (Table S5). Most species (n = 442) fell into two of four classifications: ‘Coastal’ and ‘Benthic’ or ‘Oceanic’ and ‘Pelagic’ (Fig. S4), with a few exceptions (n = 10, Table S6).

### Phylogenetic analysis

To account for the evolution of traits and phylogenetic non-independence of species, we incorporated phylogenetic trees from Stein et al.(*30*). We adopted recent taxonomic changes (Table S7) and added recently described species missing from the trees using the phytools(*61*) and ape(*62*) packages in R. In cases where multiple species were synonymized (e.g., *Cephaloscyllium maculatum* and *C. pardelotum* are synonyms of *C. fasciatum*(*63*); Table S7) we visually confirmed that the data were the same for synonymized species and selected one species entry. When adding new species (e.g., *Apristurus nakayai*(*64*)), we assigned branch lengths using the Grafen method, which accounts for shared ancestry without requiring divergence times(*65*). New species were added to the root of the corresponding genus with the add.species.to.genus function. For genera that are not monophyletic, species were added to the most inclusive group containing members of the genus. We trimmed tree tips to match the species present in our climatic, ecological, and morphological data, and vice versa, resulting in a complete data set for 452 species (82% of all sharks). Species excluded from analysis and the reason for exclusion are available in Table S8.

In order to visualize the present phylogenetic patterns in CFAR, we used the ggtree(*66*) and ggtreeExtra(*67*) packages to visualize the phylogenetic variation in present-day CFAR, maximum body size, gill slit height, water column position, and tracking data availability(*13*, *16*).

### Mapping global patterns of dispersal potential

To visualize the geographic distribution of CFAR, we computed the mean and variation in CFAR of species present by equal-area hexagonal grid cells. Using the sf package in R(*57*), we generated cells to cover the whole ocean surface, with each cell covering ∼23,322 km^2^. We intersected all available species ranges by the hexagonal grid cells to get a list of species present in each cell. If there was any spatial overlap between a species’ range and the cell, we would consider the species as present. We used equal area cells to account for variation in species range shape, size, and accuracy, as well as meaningfully compare values between cells(*45*). From this list, we matched species names to our CFAR data to generate the mean value and variability across all species present in each cell. All spatial layers were transformed or generated with the ESRI:54034 (Cylindrical Equal Area) projection. These values were then visualized on a map using ggplot2(*68*) and with hexbin plots using the hexbin package(*69*) in R. Using the same approach, we also mapped the geographic distribution of CFAR in pelagic species, coastal species, and deepwater species separately, as well as mean logged maximum body size.

### Causal inference

We followed the causal modelling framework workflow used by other causal inference studies in ecology(*35*, *70*, *71*). First, we created an initial Directed Acyclic Graph (DAG) representing how each variable may influence or cause each other and CFAR based on ecological knowledge and our own research (Fig. S5). We then critically evaluated the DAG through discussion, resulting in two possible DAGs (Fig. S6a). Each possible DAG was created to address uncertainty regarding whether water column position drives habitat use (being pelagic drives a species to occupy offshore environments) or vice versa (being offshore drives a species to occupy a pelagic water column position).

After creating the two possible DAGs, we used the phylopath package(*72*) to identify the most plausible DAG and test conditional independencies(*71*). Since CFAR has a log-linked Gamma distribution rather than a normal, or Gaussian distribution (which is the default in phylopath), CFAR was logged prior to analysis. We found that average dissolved oxygen was too highly correlated with average temperature, causing our models to fail the d-separation test (Fig. S6b, Table S9). After excluding average dissolved oxygen, we found that habitat use (coastal or oceanic) was highly correlated with water column position (benthic or pelagic) and effectively redundant (Fig. S7, Table S10). Thus, we removed habitat use as a variable and then used the phylo_path function to evaluate our new DAG (Fig. 5, Table 1). We used the best function to obtain standardized point estimates for path coefficients from the model(*71*).

### Bayesian generalized linear multilevel models

To determine the total effect (direct and indirect paths(*78*)) of each variable on CFAR, we used DAGitty to identify adjustment sets (e.g., to estimate the total effect of gill slit height on CFAR) for our final model from phylopath (https://dagitty.net/mf7gGFe4F)(79). Based on our final causal model, each variable of interest required one adjustment set (Table 2). We ran Bayesian generalized linear multilevel models on each adjustment set using the brms package(*52*) with phylogeny as a random effect. All models were fitted with four chains each of 3,000 iterations with 1,000 warm-up iterations (4,000 iterations total) and a log-link Gamma distribution due to the zero-bounded, continuous, and skewed nature of our response variable (CFAR). Model convergence was assessed by ensuring R-hat values were equal to one and effective sample size (ESS) were greater than 1,000. All analyses were performed using R (v 4.6.1).

All data and code are available in GitHub (https://github.com/raitchison/global-shark-dispersal.git).

### Cause or consequence of CFAR

It can be difficult to determine cause versus consequence in ecology so we briefly consider two alternate possible hypotheses: diet (which we included in our initial DAG, Fig. S5) and geographic range size. Diet is perhaps better viewed as an indirect consequence of CFAR since it is most often directly predicted by the interaction of mouth morphology(*80*, *81*) and prey sizes(*82*, *83*). In addition, diet is likely a more direct consequence of body size and water column position than CFAR *per se* and thus not relevant to include while evaluating the causes of CFAR. Similarly, we did not consider geographic range size in our causal analysis because in other vertebrate taxa, such as bats and birds, wing shape is considered a predictor of geographic range size(*19*, *24*, *84*). We also demonstrate that CFAR is positively related to geographic range size (see “Evaluating swimming speed and geographic range” section).

## Acknowledgements

RMA would like to thank Hannah Watkins for Bayesian insight and guidance as well as the “DAG Support Group” (Jennifer Bigman, Simon Dedman, Chris Mull, and Sophie Tattrie). RMA would also like to thank Amanda Arnold, Nina Faure Beaulieu, Owen Kui, Logan Vuorensivu, Emily Warren, and Tony Williams for their constructive comments on the manuscript.

## Author contributions

Conceptualization: RMA, NKD

Data Curation: RMA, WJV, JHM, NKD

Investigation: RMA, WJV, JHM, NKD

Methodology: RMA, WJV, JHM, NKD

Visualization: RMA

Supervision: RMA, NKD

Writing—original draft: RMA

Writing—review & editing: RMA, WJV, JHM, NKD

## Data, code, and materials availability

All data, code, and materials used are available in GitHub: https://github.com/raitchison/global-shark-dispersal

