## Supplementary Material for "From wings to tails: a general framework for inferring dispersal potential from morphology"

**Contents**

Supplementary Analysis 1: Tagging data analysis. Fig S1, Table S1.

Table S2. Bayesian model summary for measures of dispersal (swimming speed and geographic range size).

Supplementary Analysis 2: Categorical variable analysis. Figs S2 and S3, Tables S3 and S4.

Fig S4. CFAR binned by habitat use and water column position category.

Table S5. List of species with water column position classification changes per the IUCN Habitat and Ecology data.

Table S6. The ten species that are not ‘Coastal’ and ‘Benthic’ or ‘Oceanic’ and ‘Pelagic’.

Table S7. Taxonomic changes adopted in phylogenetic trees, including spelling corrections, based on Eschmeyer’s Catalog of Fishes.

Table S8. Species excluded from analysis and the reason for exclusion.

Fig S5. Initial proposed DAG.

Supplementary Analysis 3: Phylogenetic path analysis. Figs S6 and S7, Tables S9 and S10.

Fig S8. Global patterns in mean logged maximum body size.

Table S11. Model parameters for the final model evaluated in phylopath.

Supplementary Analysis 1: Tagging data analysis. Evaluating whether tagging data is biased. Code available in “1_taggingAnalysis.Rmd”.

Table S1. Chi-squared test results for each categorical axis (Water Column Position, Family, and IUCN Category).

| Axis | Chi sq | Degrees of Freedom | p-value |
|---|---|---|---|
| Water Column Position | 51.03611 | 2 | 8.27e-12 |
| Family | 177.0171 | NA (simulated) | 1.99e-4 |
| IUCN Category | 25.76682 | 5 | 9.90e-05 |

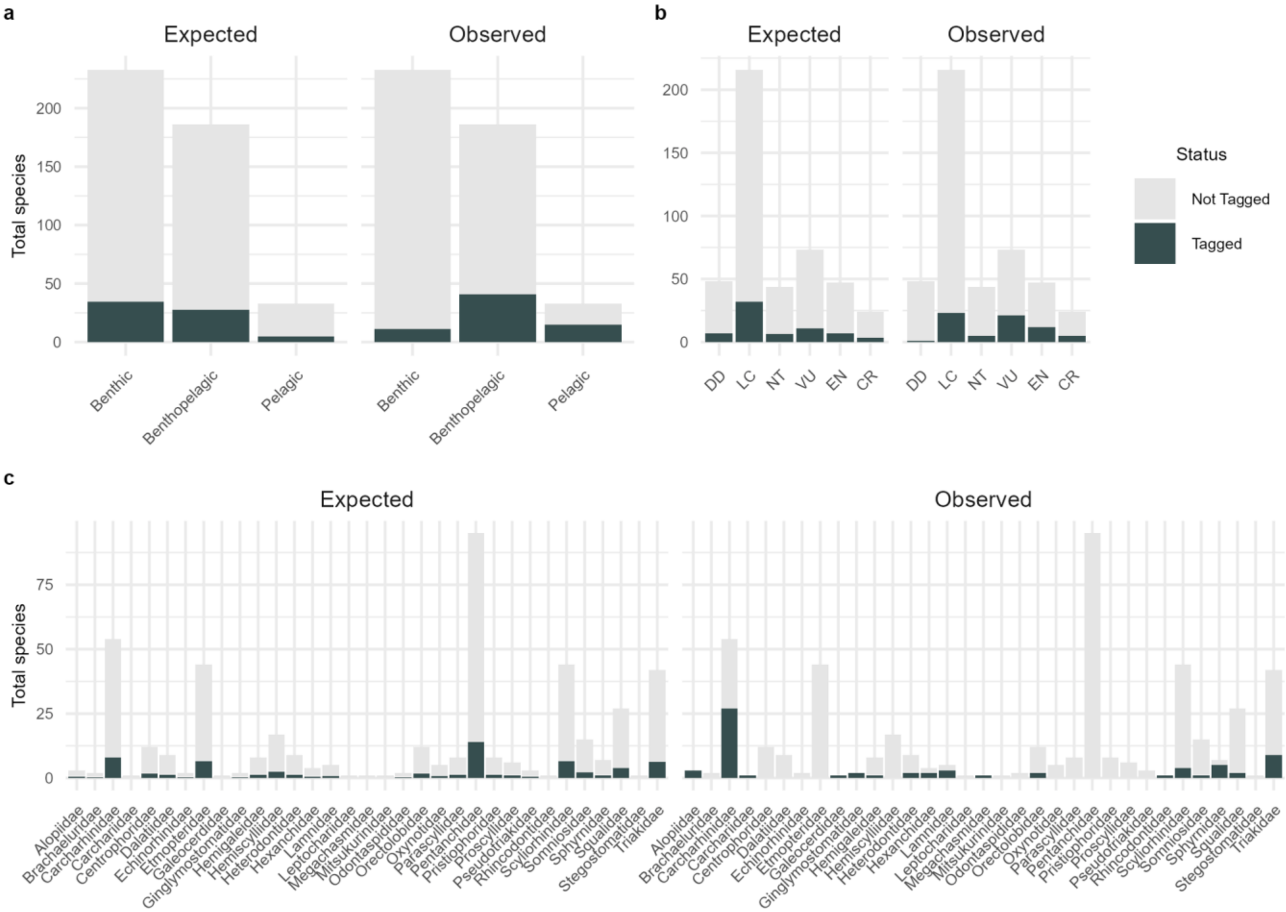


Fig S1. Expected versus observed number of species tagged. a. By water column position, b. IUCN category, c. family.

Table S2. Bayesian model summary for measures of dispersal (swimming speed and geographic range size).

| | **Estimate** | **Estimated Error** | **95% Credible Interval** |
|---|---|---|---|
| Swimming Speed ~ CFAR | | | |
| Intercept | 0.15 | 0.15 | -0.18–0.43 |
| CFAR | 0.09 | 0.05 | -0.01–0.19 |
| Geographic Range Size ~ CFAR | | | |
| Intercept | 13.17 | 1.51 | 10.16–16.20 |
| CFAR | 0.57 | 0.14 | 0.29–0.85 |

Supplementary Analysis 2: Categorical variable analysis. Reducing categorical variables (water column position and habitat use) to binary categories using phylogenetic generalized least squared (PGLS) models. Code available in “2_categoricalAnalysis.Rmd”.

*Water Column Position - benthic, benthopelagic, and pelagic*

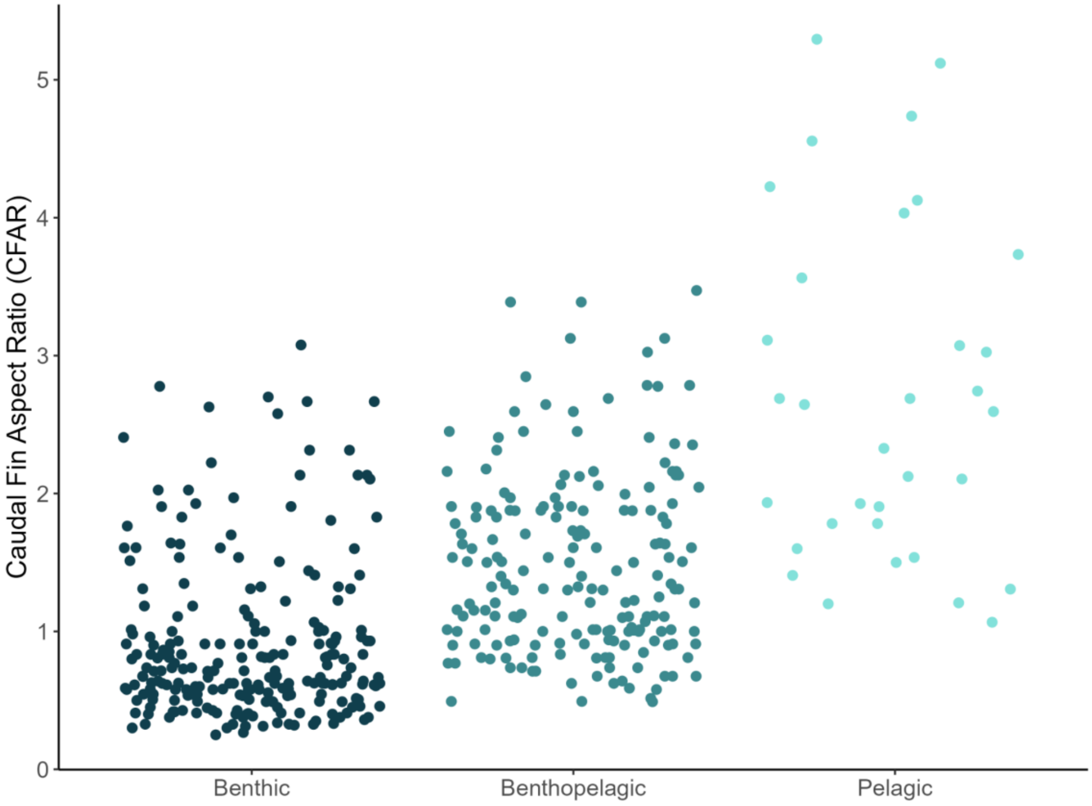


Fig S2. CFAR by water column position category. Each dot represents one species. Water column position (benthic, benthopelagic, and pelagic) were obtained from Pimiento et al. (2023) and the IUCN Red List ‘Habitat & Ecology’ narratives (see also Table S3).

Table S3. AIC values for water column position models.

| **Models** | **Degrees of Freedom** | **AIC** |
|---|---|---|
| Benthic, Benthopelagic, Pelagic | 4 | 962.5147 |
| Benthopelagic to Pelagic | 3 | 1058.7490 |
| Benthopelagic to Benthic | 3 | 964.3918 |

*Habitat Use - deepwater, coastal, and oceanic*

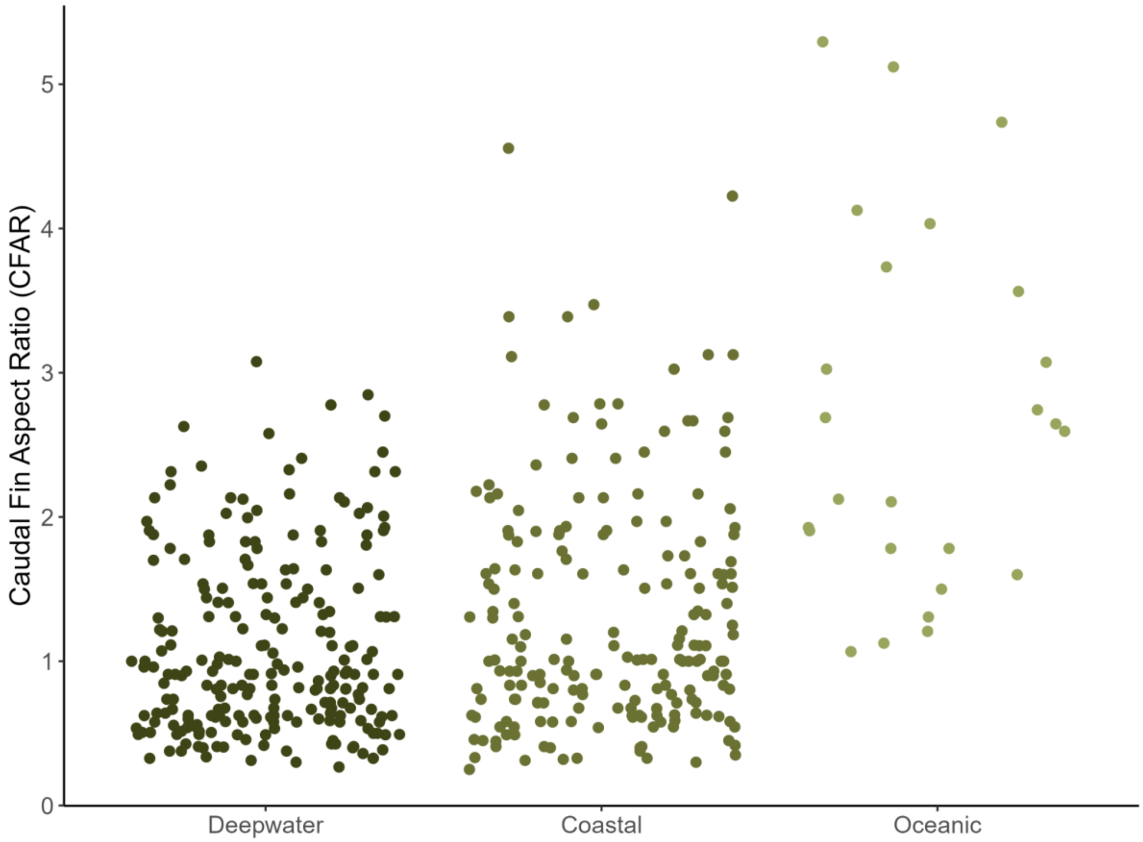


Fig S3. CFAR by habitat use category. Each dot represents one species. Habitat use (deepwater, coastal, and oceanic) were obtained from the IUCN Red List ‘Habitat & Ecology’ narratives. To avoid confusion with the water column position category “pelagic”, the habitat use category “pelagic” was renamed to “oceanic”.

Table S4. AIC values for habitat use models.

| Models | Degrees of Freedom | AIC |
|---|---|---|
| Coastal, Deepwater, Oceanic | 4 | 1034.289 |
| Deepwater to Coastal | 3 | 1039.464 |
| Deepwater to Oceanic | 3 | 1042.269 |

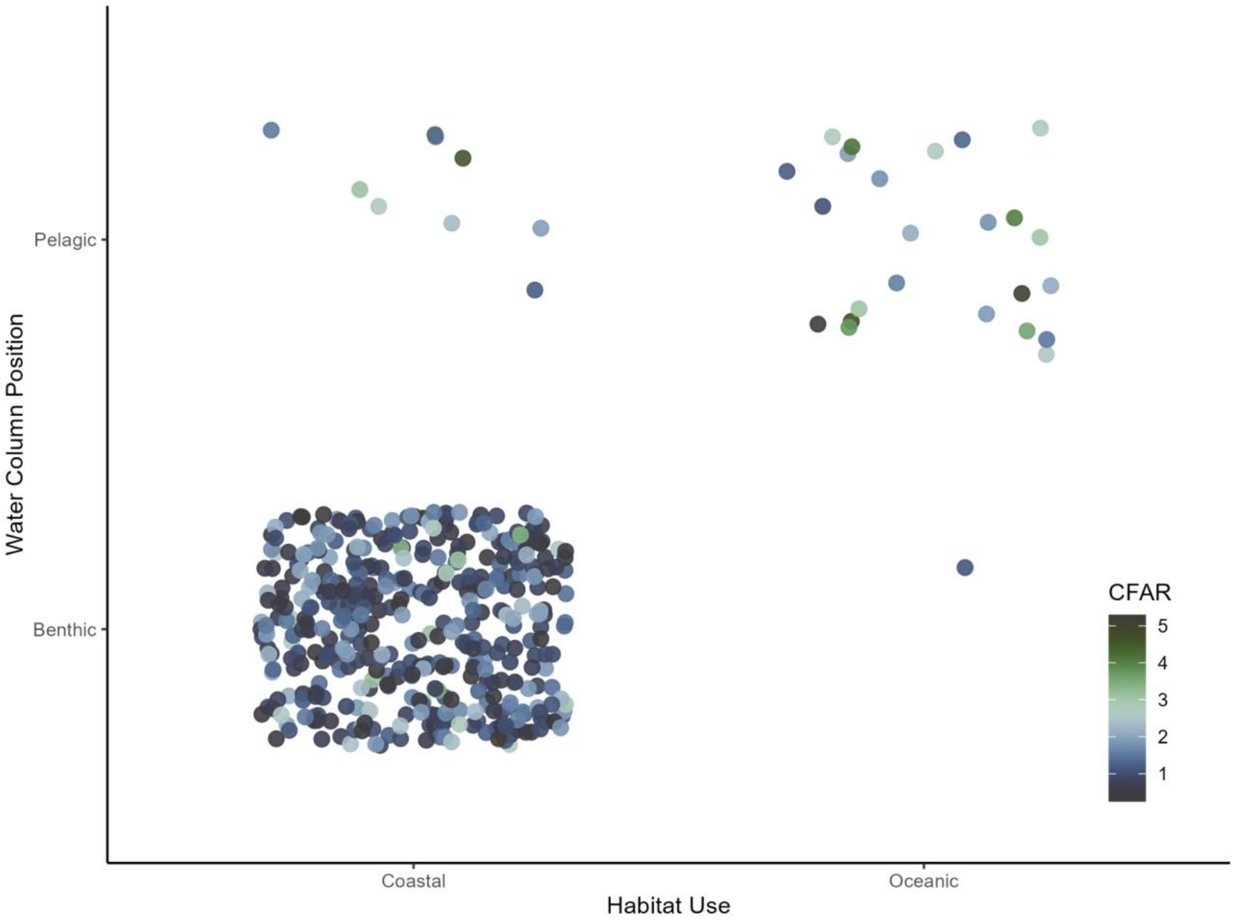


Fig S4. CFAR binned by habitat use and water column position category. Each dot represents one species. Blue colors indicate lower values and green colors indicate higher values. To avoid confusion with the water column position category “pelagic”, the habitat use category “pelagic” was renamed to “oceanic”.

Table S5. List of species with water column position classification changes per the IUCN Habitat and Ecology narratives.

| Species | Pimiento et al. 2023 Position | Re-classification based on IUCN |
|---|---|---|
| *Carcharhinus galapagensis* | Benthic | Pelagic |
| *Carcharhinus macloti* | Pelagic | Benthic |
| *Carcharhinus signatus* | Benthic | Pelagic |
| *Carcharhinus tilstoni* | Pelagic | Benthic |
| *Glyphis garricki* | Pelagic | Benthic |
| *Scymnodalatias garricki* | Pelagic | Benthic |
| *Squalus altipinnis* | Pelagic | Benthic |
| *Squalus bucephalus* | Pelagic | Benthic |
| *Squalus crassispinus* | Pelagic | Benthic |
| *Squalus edmundsi* | Pelagic | Benthic |
| *Squalus formosus* | Pelagic | Benthic |
| *Squalus nasutus* | Pelagic | Benthic |
| *Squalus notocaudatus* | Pelagic | Benthic |
| *Squalus raoulensis* | Pelagic | Benthic |

Table S6. The ten species that are not ‘Coastal’ and ‘Benthic’ or ‘Oceanic’ and ‘Pelagic’.

| **Species** | **Classification** | **Details** |
|---|---|---|
| *Carcharhinus amblyrhynchoides* | Coastal and Pelagic | Occurs inshore, from the surface to a depth of ~ 50 m. |
| *Carcharhinus brachyurus* | Coastal and Pelagic | Occurs on continental shelves from inshore to 145 m depth. |
| *Carcharhinus brevipinna* | Coastal and Pelagic | Occurs on continental and insular shelves, from the surface to a depth of 200 m. |
| *Carcharhinus humani* | Coastal and Pelagic | Likely occurs inshore and offshore, at depths up to 43 m. |
| *Carcharhinus tilstoni* | Coastal and Pelagic | Occurs in coastal waters to the continental shelf, |
| *Euprotomicroides zantedeschia* | Coastal and Pelagic | A data poor species that occurs from depths of 195 m to 641 m. |
| *Euprotomicrus bispinatus* | Coastal and Pelagic | Assumed to be circumglobal, occurs from the surface to a depth of 1,500 m. |
| *Isistius brasiliensis* | Coastal and Pelagic | Occurs from the surface to a depth of 3,500 m, and migrates vertically. |
| *Mitsukurina owstoni* | Oceanic and Benthic | Occurs well offshore, including seamounts, often at depths > 200 m. |
| *Scymnodalatias albicauda* | Coastal and Pelagic | Occurs from depths of 140 m to 512 m, and likely migrates vertically. |

Table S7. Taxonomic changes adopted in phylogenetic trees, including spelling corrections, based on Eschmeyer's Catalog of Fishes (Fricke et al. 2026; https://researcharchive.calacademy.org/research/ichthyology/catalog/fishcatmain.asp).

| Old Taxonomy | New Taxonomy |
|---|---|
| *Carcharhinus perezi* | *Carcharhinus perezii* |
| *Centrophorus lusitanicus* | *Centrophorus granulosus* |
| *Centrophorus zeehaani* | *Centrophorus uyato* |
| *Centroscymnus macracanthus* | *Scymnodon macracanthus* |
| *Centroscymnus crepidater* | *Centroselachus crepidater* |
| *Cephaloscyllium maculatum* | *Cephaloscyllium fasciatum* |
| *Cephaloscyllium pardelotum* | *Cephaloscyllium fasciatum* |
| *Chiloscyllium hasselti* | *Chiloscyllium hasseltii* |
| *Deania hystricosa* | *Deania calceus* |
| *Deania calcea* | *Deania calceus* |
| *Glyphis fowlerae* | *Glyphis gangeticus* |
| *Glyphis siamensis* | *Glyphis gangeticus* |
| *Isistius labialis* | *Isistius brasiliensis* |
| *Miroscyllium sheikoi* | *Etmopterus sheikoi* |
| *Mustelus mangalorensis* | *Iago mangalorensis* |
| *Mustelus walkeri* | *Mustelus antarcticus* |
| *Parmaturus albimarginatus* | *Dichichthys albimarginatus* |
| *Parmaturus bigus* | *Dichichthys bigus* |
| *Parmaturus melanobranchus* | *Dichichthys melanobranchus* |
| *Parmaturus nigripalatum* | *Dichichthys nigripalatum* |
| *Proscyllium venustum* | *Proscyllium habereri* |
| *Proscymnodon macracanthus* | *Scymnodon macracanthus* |
| *Proscymnodon plunketi* | *Scymnodon macracanthus* |
| *Scyliorhinus besnardi* | *Scyliorhinus haeckelii* |

| | |
|---|---|
| *Scyliorhinus tokubee* | *Scyliorhinus torazame* |
| *Scymnodon plunketi* | *Scymnodon macracanthus* |
| *Stegostoma fasciatum* | *Stegostoma tigrinum* |
| *Zameus ichiharai* | *Scymnodon ichiharai* |

Table S8. Species excluded from analysis and the reason for exclusion.

| Species | Reason for exclusion |
|---|---|
| *Akheilos suwartanai* | No water column position data |
| *Apristurus internatus* | No range data |
| *Apristurus micropterygeus* | No range data |
| *Apristurus sibogae* | No range data |
| *Bythaelurus alcockii* | Only “presence unknown” range data |
| *Bythaelurus clevai* | No temperate data |
| *Bythaelurus incanus* | No climate data |
| *Carcharhinus hemiodon* | Only "possibly extant" range data |
| *Carcharhinus wheeleri* | Taxonomic status uncertain, missing all data |
| *Cephaloscyllium formosanum* | No range data |
| *Cetorhinus maximus* | No gill slit data |
| *Chlamydoselachus africana* | No gill slit data |
| *Chlamydoselachus anguineus* | No gill slit data |
| *Deania calceus* | Taxonomic status uncertain, missing all data |
| *Etmopterus granulosus* | Missing shape file |
| *Etmopterus schmidti* | Taxonomic status uncertain, missing all data |
| *Eusphyra blochii* | No climate data |
| *Glyphis sp.1* | Invalid species, uncertain of identity |
| *Gogolia filewoodi* | No range data |
| *Gollum suluensis* | No range data |
| *Hexanchus vitulus* | No gill slit data |
| *Iago mangalorensis* | Taxonomic status uncertain, missing water column position data |
| *Mustelus andamanensis* | No CFAR data |
| *Mustelus antarcticus* | Taxonomic uncertainty in data (many synonyms) |
| *Mustelus mangalorensis* | No range data |

| | |
|---|---|
| *Parmaturus sp.* | Invalid species, uncertain of identity |
| *Parmaturus campechiensis* | No range data |
| *Pseudocarcharias kamoharai* | No climate data |
| *Pseudoginglymostoma brevicaudatum* | No climate data |
| *Scyliorhinus duhamelii* | No water column position data |
| *Scyliorhinus garmani* | No range data |
| *Scyliorhinus hachijoensis* | No CFAR data |
| *Scymnodon macracanthus* | Taxonomic uncertainty in data (many synonyms) |
| *Sphyrna gilberti* | No range data |
| *Squalus shiraii* | No CFAR data |
| *Squatina aculeata* | No gill slit data |
| *Squatina africana* | No gill slit data |
| *Squatina albipunctata* | No gill slit data |
| *Squatina argentina* | No gill slit data |
| *Squatina armata* | No gill slit data |
| *Squatina australis* | No gill slit data |
| *Squatina caillieti* | Single point range data |
| *Squatina californica* | No gill slit data |
| *Squatina david* | No gill slit data |
| *Squatina dumeril* | No gill slit data |
| *Squatina formosa* | No gill slit data |
| *Squatina guggenheim* | No gill slit data |
| *Squatina japonica* | No gill slit data |
| *Squatina legnota* | No range data |
| *Squatina mapama* | No CFAR data |
| *Squatina nebulosa* | No gill slit data |
| *Squatina oculata* | No gill slit data |

| | |
|---|---|
| *Squatina occulta* | No gill slit data |
| *Squatina pseudocellata* | No gill slit data |
| *Squatina squatina* | No gill slit data |
| *Squatina tergocellata* | No gill slit data |
| *Squatina tergocellatoides* | No gill slit data |
| *Squatina varii* | No gill slit data |

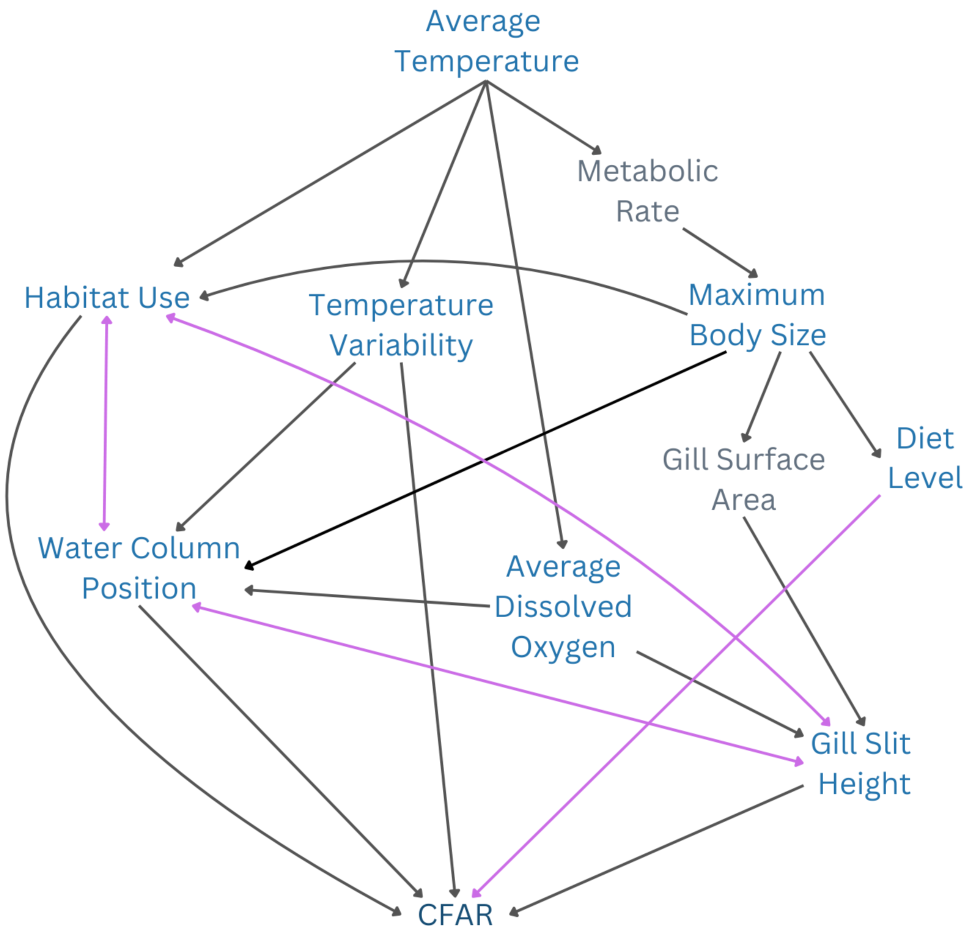


Fig S5. Initial proposed directed acyclic graph (DAG). Grey variables indicate variables where data are largely not available for sharks. Purple arrows indicate uncertainty or competing hypotheses (e.g. whether including diet is appropriate and whether water column position drives habitat use or vice versa).

Supplementary Analysis 3: Phylogenetic path analysis. Evaluating DAGs that contain dissolved oxygen and habitat use in `phylopath`. Code available in “3_phylopath.Rmd”.

*Dissolved Oxygen models*

Table S9. Model parameters for models containing average dissolved oxygen (DO). In `phylopath`, significant p-values indicate the model should be rejected.

| Model | k | q | C | p | CICc | delta_CICc | l | w |
|---|---|---|---|---|---|---|---|---|
| DO_One | 8 | 20 | 54.92 | 3.67e-06 | 96.91 | 0 | 1 | 0.99 |
| DO_Two | 8 | 20 | 63.59 | 1.29e-07 | 105.58 | 8.67 | 0.01 | 0.01 |

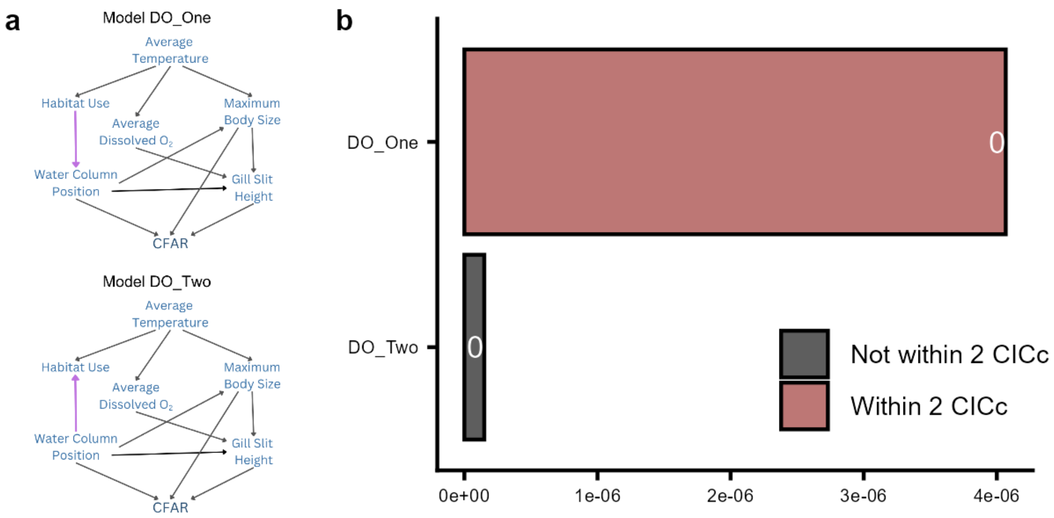


Fig S6. Phylogenetic path analysis of all proposed drivers of caudal fin aspect ratio (CFAR). a. Initial proposed DAGs, including dissolved oxygen. Purple arrows indicate competing hypotheses (tested relationships). b. Model results showing p-values (bar labels). Significance indicates rejection.

*Habitat Use models*

Table S10. Model parameters for models containing habitat use. In `phylopath`, significant p-values indicate the model should be rejected.

| Model | k | q | C | p | CICc | delta_CICc | l | w |
|---|---|---|---|---|---|---|---|---|
| Hab_One | 5 | 16 | 9.77 | 0.46 | 43.02 | 0 | 1 | 0.94 |
| Hab_Two | 5 | 16 | 15.17 | 0.13 | 48.42 | 5.40 | 0.07 | 0.06 |

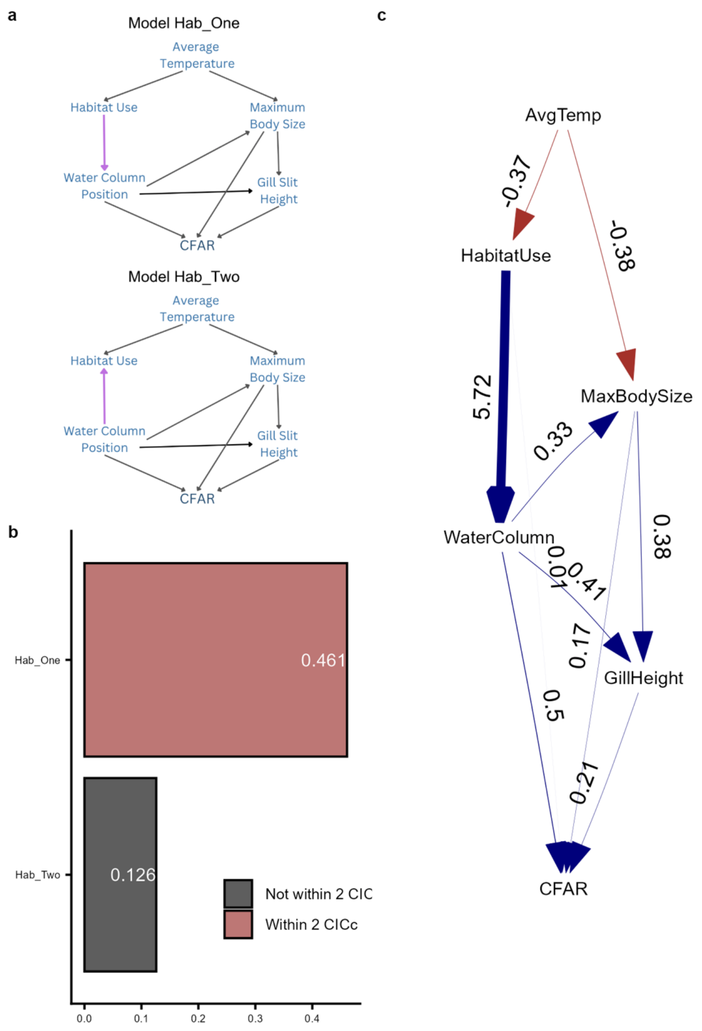


Fig S7. Phylogenetic path analysis of drivers of CFAR without dissolved oxygen. a. Two proposed DAGs. Purple arrows indicate competing hypotheses (tested relationships). b. Model results showing p-values (bar labels). Significance indicates rejection. Color indicates whether the models are within 2 CICc (red) or not (grey). c. Best model that includes habitat use with a very large path coefficient (5.72) from water column position to habitat use. Color indicates direction: positive (blue, e.g. MaxBodySize to GillSlitHeight) and negative (red, e.g. AvgTemp to MaxBodySize).

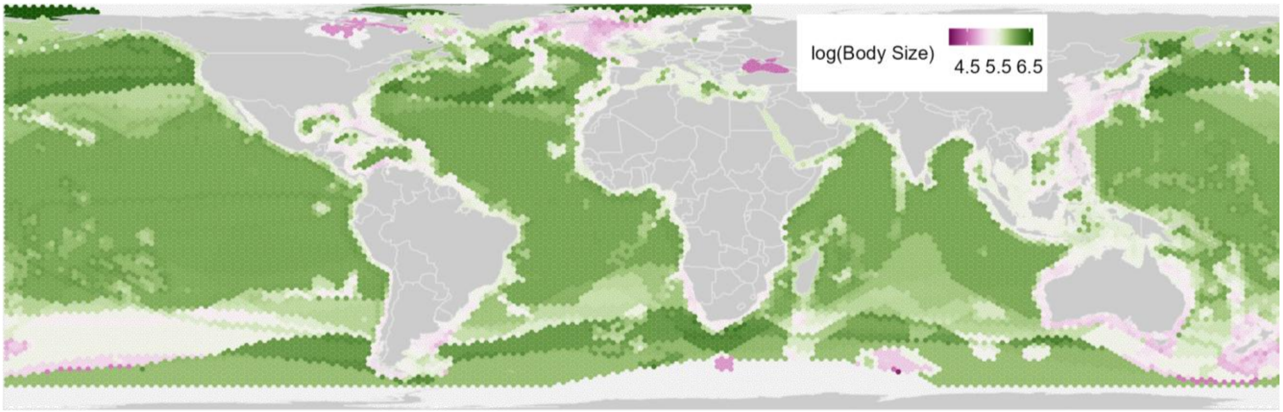


Fig S8. Global pattern in mean logged maximum body size. Pink colors indicate lower values and green colors indicate higher values. Each cell represents approximately 23,322 $km^2$.

Additional results from phylogenetic path analysis.

Table S11. Model parameters for the final model evaluated in `phylopath`.

| k | q | C | p | CICc | delta_CICc | l | w |
|---|---|---|---|---|---|---|---|
| 3 | 12 | 5.26 | 0.51 | 29.97 | 0 | 1 | 1 |
